# Thermal stress triggers productive viral infection of a key coral reef symbiont

**DOI:** 10.1101/2021.03.17.435810

**Authors:** Carsten GB Grupstra, Lauren I Howe-Kerr, Alex J Veglia, Reb L Bryant, Samantha R Coy, Patricia L Blackwelder, Adrienne MS Correa

**Author notes:** These authors contributed equally to this work (Lauren I Howe-Kerr, Alex J Veglia). **Competing Interests statement:** The authors declare that they have no competing interests.

## Abstract

Climate change-driven ocean warming is increasing the frequency and severity of bleaching events, in which corals appear whitened after losing their dinoflagellate endosymbionts (family Symbiodiniaceae). Viral infections of Symbiodiniaceae may contribute to some bleaching signs, but little empirical evidence exists to support this hypothesis. We present the first temporal analysis of a lineage of Symbiodiniaceae-infecting positive-sense single-stranded RNA viruses (‘dinoRNAVs’) in coral colonies, which were exposed to a 5-day heat treatment (+2.1°C). A total of 124 dinoRNAV major capsid protein gene ‘aminotypes’ (unique amino acid sequences) were detected from five colonies of two closely related *Pocillopora-Cladocopium* (coral-symbiont) combinations in the experiment; most dinoRNAV aminotypes were shared between the two coral-symbiont combinations (64%) and among multiple colonies (82%). Throughout the experiment, seventeen dinoRNAV aminotypes were found only in heat-treated fragments, and 22 aminotypes were detected at higher relative abundances in heat-treated fragments. DinoRNAVs in fragments of some colonies exhibited higher alpha diversity and dispersion under heat stress. Together, these findings provide the first empirical evidence that exposure to high temperatures triggers some dinoRNAVs to switch from a persistent to a productive infection mode within heat-stressed corals. Over extended time frames, we hypothesize that cumulative dinoRNAV production in the *Pocillopora-Cladocopium* system could affect colony symbiotic status, for example, by decreasing Symbiodiniaceae densities within corals. This study sets the stage for reef-scale investigations of dinoRNAV dynamics during bleaching events.

## Introduction

Warming seas, driven by climate change, are increasingly causing bleaching events: mass losses of endosymbiotic dinoflagellates (family Symbiodiniaceae) from corals and other invertebrate hosts. Bleaching events often result in coral mortality and are contributing to the degradation of reef ecosystems globally [1, 2]. Viruses, which are diverse and abundant on coral colonies [3–7], are hypothesized to contribute to some coral bleaching signs by lysing Symbiodiniaceae cells (e.g., [8–10]). Alternatively, viral shifts in conjunction with bleaching-associated stressors (e.g., [11– 14]) could merely be correlated with bleaching signs or constitute opportunistic secondary infections [15, 16]. Beyond bleaching, viruses may influence colony health by altering the function of resident microbial symbionts or coral tissues (e.g., [17–23]). Although various roles for viruses in coral bleaching, disease, and function have been hypothesized [23, 24], thus far, these roles have been difficult to test empirically.

Symbiodiniaceae are putative target hosts of DNA and RNA viruses (reviewed in [6, 7]), including the ‘dinoRNAVs’, a group of dinoflagellate-infecting positive-sense single-stranded RNA viruses. Although dinoRNAVs have yet to be isolated, stably propagated, and fully characterized, they have been detected in transcriptomes from Symbiodiniaceae cultures, as well as in viral particles isolated from Atlantic and Pacific corals spanning 6 genera (Table 1). These associations suggest that dinoRNAVs are prevalent as exogenous, persistent infections in Symbiodiniaceae cells [8–10, 25,26]. Furthermore, in previous studies, increased dinoRNAV detection [25], and changes in dinoRNAV and host anti-viral transcript expression in a thermosensitive symbiont population only [27], suggest that under stressful conditions, some dinoRNAVs may switch to a more productive replication mode that could culminate in host lysis.

**Table 1.**
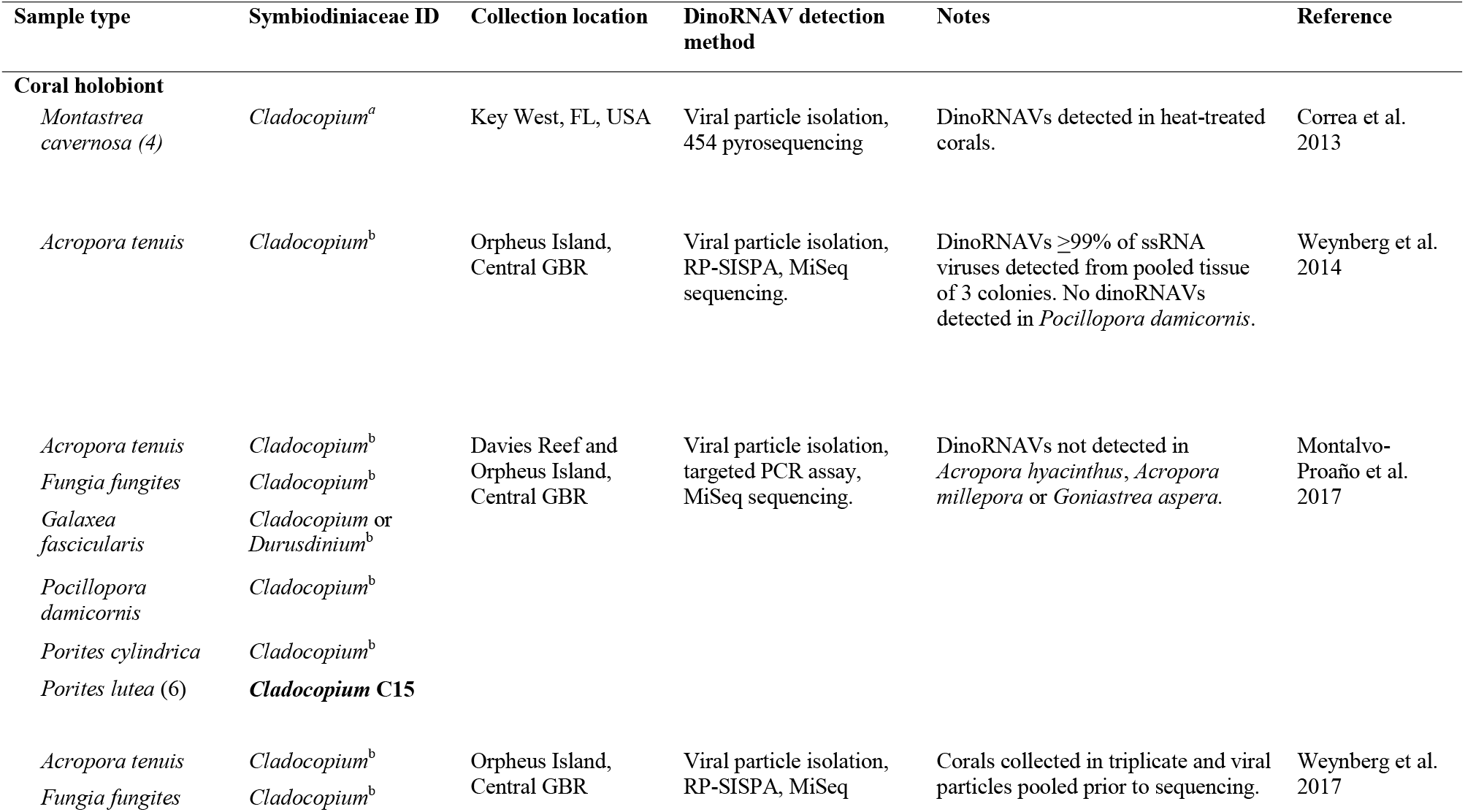
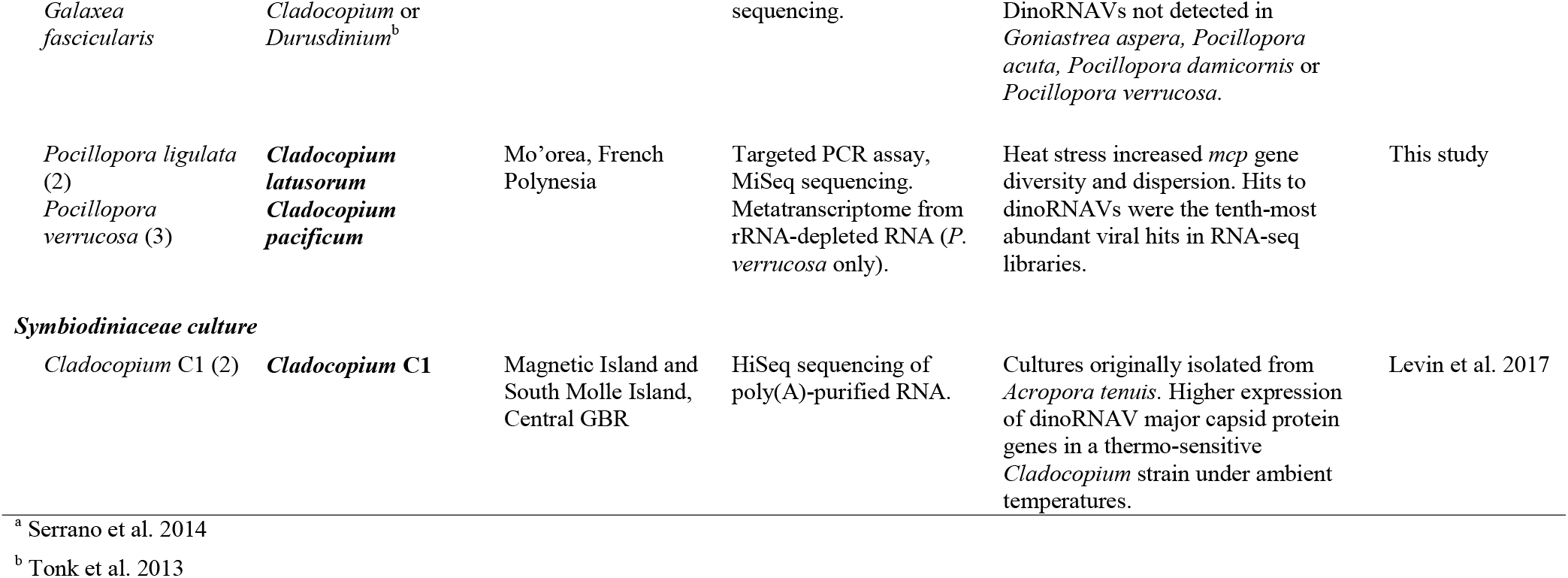
Summary of studies in which Symbiodiniaceae-infecting dinoflagellate RNA virus (dinoRNAV) genes have been recovered from coral colonies or Symbiodiniaceae cultures. Samples sizes are n=1 unless indicated otherwise in parentheses. **Bolded** Symbiodiniaceae IDs were directly characterized in a given study. All other Symbiodiniaceae IDs indicate the Symbiodiniaceae genus (or genera) typically reported from a given coral species based on published literature.

This infection strategy has recently been identified in other algal host-virus systems (e.g., coccolithophore Emiliana huxleyi-EhV system, [28]).

Heterocapsa circularisquama RNA virus (HcRNAV), which infects free-living dinoflagellates, is among the closest known relatives to Symbiodiniaceae-infecting dinoRNAVs [25, 29]. HcRNAV undergoes a putatively strictly lytic replication cycle following a latent period of 24-48 hours in laboratory experiments; during this latent period, the host is infected but not yet lysed and viruses are not yet released [30]. Based on this, we hypothesized that shifts by Symbiodiniaceae-infecting viruses to a more productive replication mode might also be detectable within the first few days of exposure to stress [13, 25, 27]. Examining viral dynamics within individual coral colonies at the onset of thermal stress can help clarify whether viral infections of Symbiodiniaceae contribute to some coral bleaching signs.

With this aim, we quantified, for the first time, the temporal dynamics of dinoRNAVs within coral colonies exposed to control and heat stress conditions. DinoRNAV diversity was then characterized from heat-treated and control fragments of each colony via amplicon sequencing of the major capsid protein (*mcp*) gene. The *mcp* gene was targeted to facilitate comparisons with previous dinoRNAV detections (Table 1, [29]) and because the *mcp* gene resolved ecologically distinct strains in the related *H. circularisquama*-HcRNAV system [30, 31]. We hypothesized that: (1) dinoRNAVs are present in a majority of Mo’orean pocilloporid colonies; and (2) dinoRNAV richness increases and composition shifts within 48 hours of exposure to thermal stress. By analyzing dinoRNAV diversity at the amino acid level, this study partially circumvented methodological challenges arising from the high mutation rates and genetic diversity of single-stranded RNA viruses [32–34], which have previously made it difficult to quantify RNA viral dynamics at ecologically relevant scales [35]. As research on reef-associated dinoRNAVs continues to progress, the aminotypes presented here may eventually merit further collapse into ‘quasispecies’—heterogeneous mixtures of related genomes [35–38]—a common approach for conceptualizing diversity within RNA virus populations.

## Materials and methods

### Experimental design

We conducted a replicated aquarium experiment (two treatments; four aquaria per treatment) in which fragments from five colonies of the morphologically cryptic stony coral *Pocillopora* species complex [39], harboring *Cladocopium* spp. symbionts, were exposed to control conditions (ambient reef water; 28.2°C) or a +2.1°C heat treatment (summer bleaching temperatures; 30.3°C) for 5 days (See Figure S1 and Supplementary Methods, [40, 41]). At the start of the experiment (t_(h)_ = 0), all fragments were photographed with a Coral-Watch Health Monitoring Chart [42] in the frame, and one fragment per colony in the control aquaria (5 fragments in total) were preserved as initial controls. At five time points (t_(h)_ = 4, 12, 24, 72 and 108 h), all fragments were photographed and visually inspected for signs of stress (e.g., excessive mucus production), lesions and/or paling. To characterize the effect of the heat treatment on Symbiodiniaceae cell densities, we compared brightness values—a proxy for Symbiodiniaceae chlorophyll concentrations, and a metric of coral bleaching status—from standardized photographs of each coral fragment at the start of the experiment and at the time of preservation (See Supplementary Methods, [42, 43]). We tested for changes to brightness values using linear mixed effects models (LMM) by including an interaction between treatment and time, and including colony ID as a random effect.

A control and a heat-stressed fragment per colony were preserved at each time point (generating 10 fragments per time point). DNA and RNA were extracted from each sample (which included coral animal tissue, Symbiodiniaceae cells and viruses) using a ZymoBIOMICS DNA/RNA Miniprep Kit (Zymo Research, Irvine, CA, USA) with an additional enzyme digestion step to improve viral RNA yields.

Coral and symbiont species were delineated from extracted DNA as described in [44]. Briefly, coral species were identified by sequencing the *Pocillopora*-specific mitochondrial open reading frame (*mt-ORF*) gene region [44–48]; symbiont species were identified by sequencing the D1/D2 domain of the large ribosomal subunit (LSU rRNA gene) and the non-coding plastid minicircle (psbAncr; [44, 49, 50]). This analysis revealed that colonies 1, 4 and 5 were *Pocillopora verrucosa* corals harboring *Cladocopium pacificum* symbionts, and colonies 2 and 3 were *Pocillopora ligulata* containing *Cladocopium latusorum* symbionts [44].

### Sequencing of dinoRNAV gene amplicons, bioinformatics processing, and phylogenetic analysis

The dinoRNAV major capsid protein (*mcp*) gene was amplified from cDNA libraries (generated from extracted RNA) using a nested PCR protocol with degenerate primers [29]; cleaned and normalized libraries were sequenced on the Illumina MiSeq platform using PE300 v3 chemistry. To rule out the possibility that the sequenced gene fragments were endogenous viral elements (EVEs) integrated into Symbiodiniaceae genomes, we attempted to amplify the *mcp* gene from DNA extracted from t_(h)_ = 0 samples. No bands were detectable on agarose gels, indicating that the dinoRNAV *mcp* sequences detected in this study are most parsimoniously interpreted as exogenous infections, and not EVEs, of Mo’orean *C. latusorum* or *C. pacificum*. Processing and analysis of all raw dinoRNAV *mcp* gene reads were conducted using the program vAMPirus (v1.0; See Supplementary Methods for details; [51]). Briefly, ASVs were generated via the UNOISE [52] algorithm with vsearch v. 2.14.2 [53], and all ASVs were then translated and collapsed into ‘aminotypes’—unique amino acid sequences each differentiated by at least one amino acid. Any sequences containing stop codons were removed prior to further analysis.

An alignment was made in MUSCLE (v5) using aminotypes from this study, as well as a dataset from the Great Barrier Reef [29] that was reprocessed using the methods above, and a set of reference best BLASTx hits to the NCBI database [54, 55]. The alignment (Supplementary Data 1) was trimmed to the first column on either side that contained no gaps, and then used to determine the best model for evolution (LG+G4+I) according to ModelTest-NG [56]. A maximum-likelihood phylogeny was inferred with RAxML-NG (1000 bootstrap iterations) and rooted with HcRNAV as an outgroup.

### Statistical analyses of dinoRNAV *mcp* aminotype composition and diversity

All data processing, visualization, analysis and statistical tests were conducted in R version 4.0.2 and Vegan 2.5-6 [57] on an *mcp* aminotype counts table (Supplementary Data 2; R code doi: https://doi.org/10.1101/2021.03.17.435810). For some analyses, the dataset was rarefied to 59 837 amino acid sequences per sample. First, the overall distribution of dinoRNAV *mcp* aminotypes among cryptic coral-symbiont species and colonies was explored to determine how subsequent statistical analyses should be conducted. A PERMANOVA based on Bray-Curtis distances from square-root-transformed rarefied data was used to test for differences in overall dinoRNAV composition among species, treatments and colonies, and over time, using adonis() in vegan. We tested for an interaction between treatment and colony, with time and species as additional factors. Although dispersion differed significantly among groups (betadisper test), PERMANOVA is robust to heterogeneous dispersion if the design is balanced [58]. The PERMANOVA revealed that dinoRNAV composition differed between the coral-symbiont species (R^2^=0.21, p<0.001). However, colony ID was a stronger predictor (R^2^=0.55, p<0.001), indicating that differences in dinoRNAV composition were mainly structured at the colony level, rather than the coral-symbiont species level.

Venn diagrams were made to identify aminotypes shared among coral-symbiont species, colonies or treatments based on non-rarified data using the online tool http://bioinformatics.psb.ugent.be/webtools/Venn/ (accessed August 17^th^, 2020). For comparison, Venn diagrams were also made based on rarified data (results not shown); these Venn diagrams exhibited similar patterns but were less conservative and were therefore not included. Venn diagram analysis indicated that 64% of aminotypes (79 of 124) were shared between coral-symbiont species. Based on these preliminary explorations of the distribution of dinoRNAV *mcp* aminotypes, in all subsequent statistical tests, we analyzed all five colonies together (i.e., at the *Pocillopora-Cladocopium* level) but included coral-symbiont species as a separate factor in LMMs.

Shannon’s diversity index (H) values were calculated based on rarefied data; expected aminotype richness values were calculated using repeated random subsampling of non-rarefied data (sample size=59 837 amino acid sequences). We tested for differences in Shannon’s diversity index and aminotype richness values using LMMs with the package LME4 v1.1-23 [59]. We tested for an interaction between treatment and timepoint; by incorporating colony ID as a random effect, this analysis approaches a repeated measures test. Coral-symbiont species were included as an additional factor to test for differences between species. F-tests were used for model selection with car v3.0-8. Assumptions of normality of the residuals were assessed visually with quantile-quantile plots and Shapiro-Wilk tests; the assumption of homogeneity of variance was visually assessed using plots with residuals versus fitted values. We also tested for differences between control and heat-treated fragments at each timepoint (using colonies as replicates), as well as per colony (using time points as replicates), with ANOVAs and controlled for type 1-errors using a Bonferroni correction.

To quantify the dispersion of dinoRNAVs between treatments and over time, a non-metric multidimensional scaling (NMDS) plot was constructed based on Bray-Curtis distances from square-root-transformed rarefied data (k=2 999 iterations), and the distance to centroid for each sample was calculated. Since dinoRNAVs differed among coral colonies, we calculated centroids for each individual colony in the control and heat treatments separately (5 colonies x 2 treatments = 10 centroids) to examine the effect of heat treatment on dinoRNAV dispersion in a given coral colony. For this analysis, different timepoints were used as replicates. We tested for differences in dispersion using the same LMM approach as described above for aminotype alpha diversity, but the values were square root-transformed because they did not follow the assumption of normality of the residuals.

Lastly, we conducted a differential abundance analysis using the non-rarefied amino acid counts table with DESeq2 v1.26.0 [60]. We fitted a negative binomial model and Benjamini-Hochberg FDR-corrected Wald tests (α=0.05) were used to test for differences in taxon abundance between treatment within each colony and at each timepoint after the start of the experiment (t_(h)_= 4, 12, 24, 72, 108). We excluded all fragments sampled at timepoint 0, as well as any fragment with <10 000 reads (and its paired fragment in the other treatment) at a given timepoint (colonies 3 and 5 at timepoint t_(h)_=4, colony 3 at t_(h)_=72).

### Metatranscriptome sequencing and bioinformatics processing

As an additional test of whether the *mcp* gene amplicons generated in this study represent exogenous dinoRNAVs, we generated 5 metatranscriptomes from random timepoints and treatments for each colony of *P. verrucosa-C. pacificum* (t_(h)_=0 control for colony 1, t_(h)_=24 heat for colony 4, t_(h)_=108 control and heat for colony 5, t_(h)_=72 control for colony 5; see Supplementary Methods for more detail) and queried them for dinoRNAV-derived transcripts. All metatranscriptome reads were trimmed, quality filtered, and merged before alignment to the proteic version of the Reference Virus Database (v21) using DIAMOND BLASTx [55]. Relative abundances of hits to dinoRNAV reference sequences, and mean alignment metrics (e-values, bitscores and % identity), were calculated. All cleaned forward and reverse reads were then normalized and assembled into contigs using rnaSPAdes v3.13.0 [63]. DinoRNAV contigs were then inferred by DIAMOND BLASTx alignment (minimum bitscore50; minimum ORF length=30 aa; minimum percent identity=30) to a hybrid database containing the uniprot database and the Reference Virus Database [64]. Prodigal (v.2.6.3) was then used to predict open reading frames and protein sequences from all dinoRNAV-like contigs, and protein annotation was conducted with another round of DIAMOND BLASTp alignment to the hybrid database. Contigs with best hits to the dinoRNAV *mcp* gene, as well as *mcp* gene amplicons, were mapped to a dinoRNAV *mcp* gene assembly from cultured *Cladocopium* (species C1) [27].

### Data availability

All *mcp* gene amplicon libraries and the 5 metatranscriptomes have been deposited to the Sequence Read Archive under accession PRJNA778019.

### Code availability

Code used to generate these results is available at https://doi.org/10.1101/2021.03.17.435810.

## Results

### Coral holobiont traits

Analysis of coral and Symbiodiniaceae gene markers revealed that colonies 1, 4 and 5 were *P. verrucosa* harboring *C. pacificum*, and colonies 2 and 3 were *P. ligulata* containing *C. latusorum* [44]. Coral fragments in the control and heated aquaria remained apparently healthy throughout the experiment; no signs of stress such as paling, mucus production, or tissue sloughing were observed. Linear mixed effects models of color values did not reveal significant paling in heat-treated coral fragments (treatment F=0.10, p=0.75; timepoint F= 0.30, p=0.59; treatment*timepoint F= 0.09, p= 0.77). Lack of color change was expected; bleaching signs are generally only detectable after weeks of ecologically relevant temperature stress—even though vital molecular processes are affected within several days [65].

### DinoRNAV *mcp* gene sequencing overview

DinoRNAV *mcp* genes were detected in cDNA libraries from all 5 colonies. Amplicon sequencing of the dinoRNAV *mcp* gene resulted in 10 222 055 paired raw reads from all samples and one negative control. A total of 7 593 537 reads with a mean length of 423 bases (before trimming to 422 bases) remained after merging and quality control. Denoising resulted in 273 unique amplicon sequence variants (ASVs) across all samples, and translation revealed that 11 ASVs contained stop codons; these ASVs were removed. Translated ASVs collapsed into 124 unique amino acid sequences—’aminotypes’—that were 140 aa in length. Five control samples and the negative control were removed because they contained few (<10 000) amino acid sequences (20 reads were detected in the negative control). All remaining samples had between 50 000 and 210 000 amino acid sequences with a mean read depth of 142 810 ± 36 163 (SD).

### DinoRNA viruses in *Pocillopora-Cladocopium* colonies are similar to dinoRNAVs in other coral-symbiont species

The 124 unique dinoRNAV aminotypes (Figure 1; Table S1) detected in this study were most similar to 13 amino acid sequences translated from a published library of dinoRNAV *mcp* nucleotide sequences, generated from exogenous viral particles isolated from six coral-symbiont species via ultracentrifugation of cesium chloride gradients [29]; mean sequence similarities to these *mcp* sequences ranged from 53.4-98.4% (mean e-values ranged from 1.43·10^−71^-6.38·10^−36^). All dinoRNAV *mcp* aminotypes in this study formed a clade that is more recently derived than reference sequences such as HcRNAV, Beihai sobemo-like virus and sponge weivirus-like virus (Figure 1). The dinoRNAV aminotypes in this study may form (at least) three quasispecies (e.g., rectangles in Figure S2, *sensu* [66]). Strikingly, the majority of dinoRNAV *mcp* genes identified in our study occupy one clade; on average, sequences in this clade vary 9.8% ± 5.5 (SD) from each other (largest clade in Figure 1, largest rectangle in Figure S2; e.g., aminotypes 15, 18, and 30). Ultra-deep sequencing may be necessary to further clarify the drivers of these potential ‘mutant cloud dynamics’ in Symbiodiniaceae hosts [67]. In several branches of the tree, aminotypes from our study are most closely related to aminotypes resolved from Symbiodiniaceae in *Acropora tenuis, Favia fungites, Galaxea fascicularis, Pocillopora damicornis, Porites cylindrica*, and *Porites lutea* corals sampled from the Great Barrier Reef (Table S1; [29]). Dominant aminotypes (>1% abundance across total dataset) are present in all major branches of the tree but differ by as much as ∼60% in amino acid sequence (three branches in Figure 1, three rectangles in Figure S2; e.g. aminotypes 1, 2, and 6). Aminotype detections varied across treatment and timepoint in the experiment (shading of red and black squares in Figure 1).

**Figure 1.**
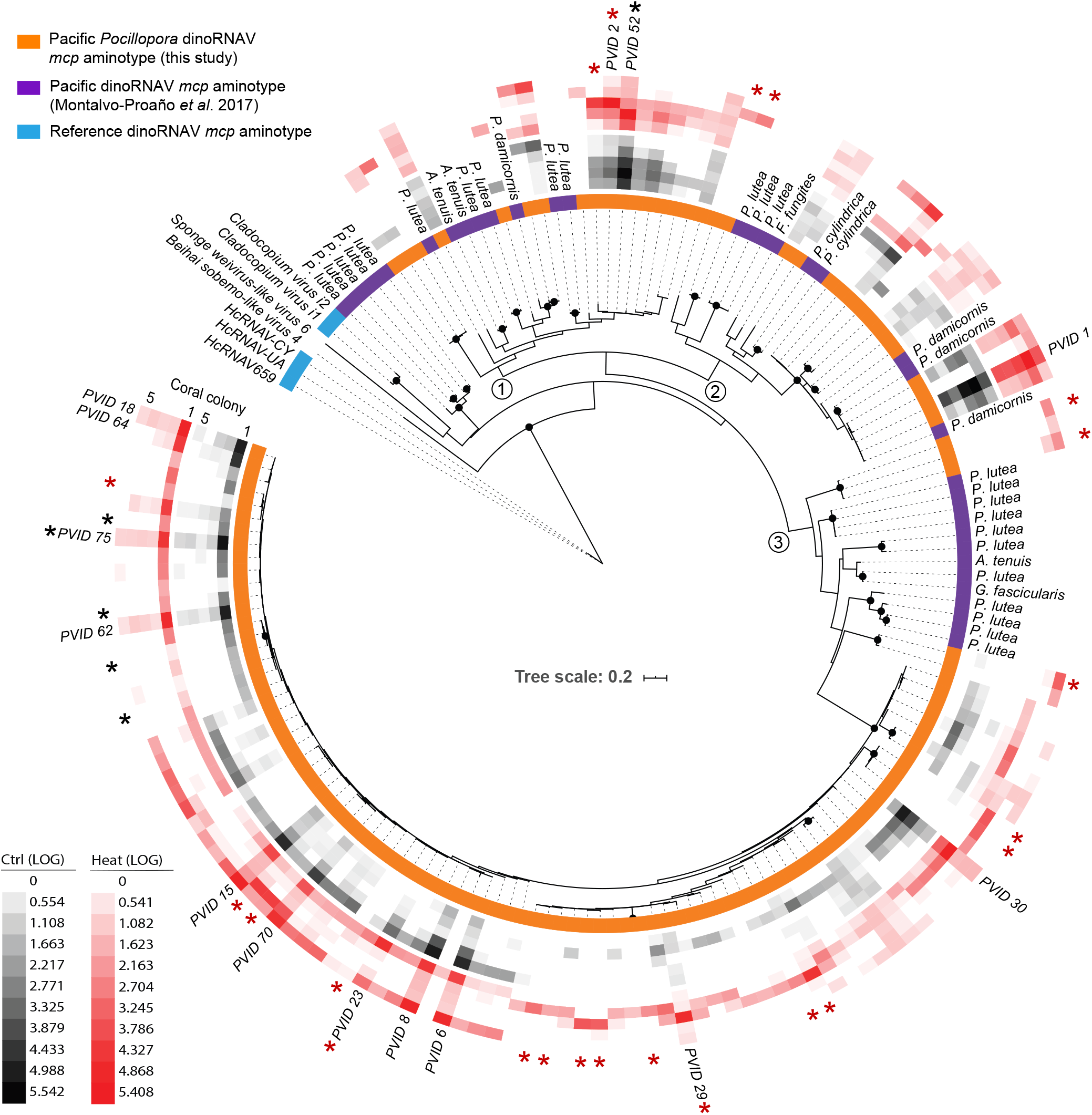
Maximum likelihood tree of major capsid protein (*mcp*) aminotypes (unique amino acid sequences) from Symbiodiniaceae-infecting dinoRNA viruses (‘dinoRNAVs’) isolated from three colonies of *Pocillopora verrucosa* and two colonies of *P. ligulata* containing the Symbiodiniaceae species *Cladocopium pacificum* and *C. latusorum*, respectively. Aminotypes recovered in this study were similar to those in a previous work that recovered dinoRNAV *mcp* sequences from viral particles isolated from six coral species via ultracentrifugation of cesium chloride gradients [29]. Three major branches of the tree are indicated with the numbers 1-3. Black dots at nodes represent bootstrap support >75%. Colors adjacent to the tree indicate the study (orange, purple) or NCBI reference (blue) for each *mcp* aminotype. Reference NCBI accession numbers: Beihai Sobemo-like Virus 4 (YP_009336877), Sponge Weivirus-like Virus 6 (ASM94037), *Cladocopium* Virus i2 (AOS87317), *Cladocopium* Virus i1 (AOG17586), HcRNAV-CY (BAE47072), HcRNAV-UA (BAE47070) and HcRNAV-659 (BAU51723). Rings with black or red squares indicate the relative, log(10)-transformed abundances of dinoRNAV aminotypes in fragments from individual coral colonies in the control or heat treatment, respectively. Aminotypes labeled with *PVID* and a numeral comprise >1% abundance of the total dataset and are included in Figure 2. Black and red asterisks indicate aminotypes that were significantly associated with control or heat treatments, respectively (see Figure 4 for details).

### DinoRNAV aminotypes differed among coral-symbiont species, colonies, and treatments

Approximately 64% of aminotypes (79 of 124) in this study were shared between the two coral-symbiont species, 19% (24) were unique to the three colonies of *P. verrucosa-C. pacificum*, and 17% (21) were unique to the two colonies of *P. ligulata-C. latusorum*. Approximately 34% of aminotypes (42) were present in all colonies, 48% (60) were shared between 2-4 colonies and 18% (22) were unique to individual coral colonies (with individual colonies containing up to 6 unique colony-specific aminotypes, Figure 2a). All 14 aminotypes that each comprised >1% reads in the total dataset (listed in Figure 2b) were detected in all five coral colonies. DinoRNAV aminotype compositions varied among individual colonies (Figure 2b); colony ID and coral-symbiont species were the most powerful predictors of viral composition (PERMANOVA: R^2^ = 0.55 and 0.21, respectively; p<0.001). DinoRNAVs responded to elevated temperatures in colony-specific ways, as indicated by a significant interaction effect between treatment and colony (PERMANOVA: R^2^=0.079, p<0.001). Treatment was also significant by itself (PERMANOVA: R^2^= 0.02, p<0.001), demonstrating a subtle but consistent response to elevated temperature across all colonies. Time was not a significant predictor (PERMANOVA: R^2^= 0.003, p=0.47; Figure 2b).

**Figure 2.**
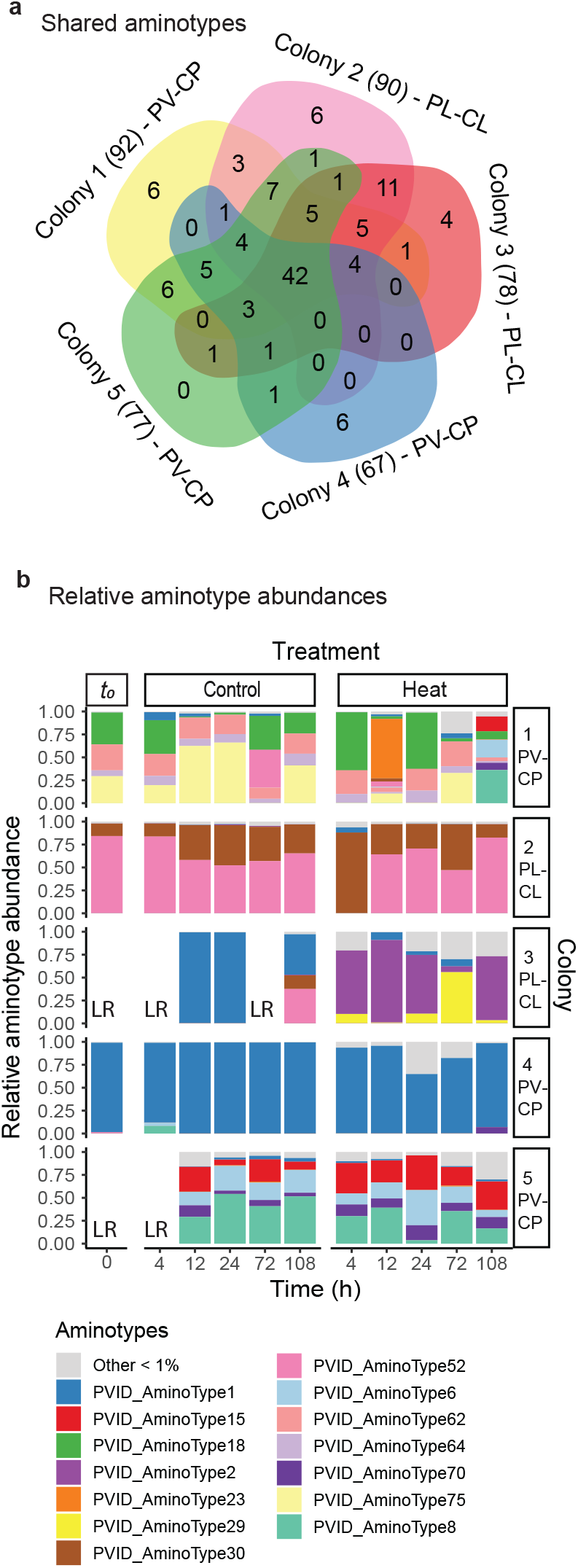
Overview of Symbiodiniaceae-infecting dinoRNAV major capsid protein (*mcp*) gene aminotypes (unique amino acid sequences) associated with the stony corals *Pocillopora verrucosa* (harboring *Cladocopium pacificum*, PV-CP) and *P. ligulata* (habroring *C. latusorum*, PL-CL). (a) **A** Venn diagram based on non-rarefied data indicates that ∼82% (102) of 124 unique aminotypes were shared between two or more coral colonies, and 64% (79) were shared between the two coral-symniont species. Data come from 8-11 fragments per colony sampled from control and heated aquaria. The total number of aminotypes detected per colony is given in parentheses. (b) The relative abundances of *mcp* aminotypes differed in fragments exposed to a 2.1 °C temperature increase (‘Heat’ treatment), compared to fragments from the same colonies exposed to ambient reef conditions (‘Control’). LR indicates “low reads” and corresponds to samples that had <10 000 reads and were excluded from the analysis. PERMANOVA results: Colony R^2^=0.55, p<0.001; Coral-symbiont species R^2^=0.21, p<0.001); Treatment R^2^=0.02, p<0.001; Treatment*Colony R^2^=0.079, p<0.001; Time R^2^=0.003, p>0.05.

### Heat treatment rapidly increased the diversity of dinoRNAV aminotypes

A total of 17 aminotypes were unique to the heat treatment; 1 aminotype was unique to the controls (Figure 3a; Table 2). Heat-specific aminotypes were observed in all colonies and ranged from 1-6 unique aminotypes per colony. Three heat-specific aminotypes were shared among multiple (2-3) coral colonies; one of these was shared between the two coral-symbiont species. The other 14 heat-specific aminotypes were not shared among colonies. A total of nine heat-specific aminotypes were unique to *P. ligulata*, whereas seven were unique to *P. verrucosa* (Table 2; Figure S2). Most heat-specific aminotypes were relatively rare; only two aminotypes comprised >1% of reads in each fragment (aminotype 25: 1.8%; aminotype 114: 1.4%; other aminotypes comprised means of 0.01-0.7% of reads in individual fragments). The single aminotype unique to controls was detected in one fragment (colony 2) after 72 hours (0.02% of reads in that fragment).

**Figure 3.**
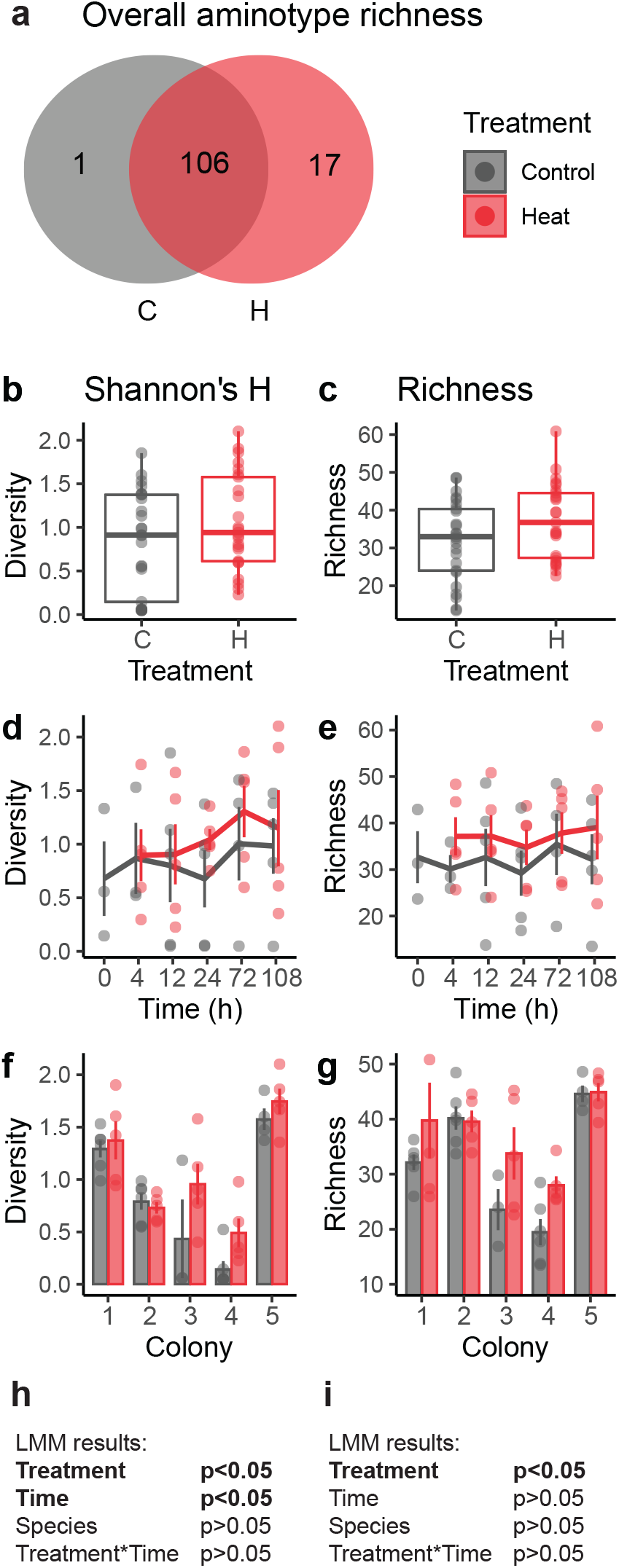
Diversity of Symbiodiniaceae-infecting dinoRNAV major capsid protein (*mcp*) gene aminotypes (unique amino acid sequences) in heat (‘Heat’, ‘H’) versus control (‘Control’, ‘C’) conditions. (a) Venn diagram of non-rarified aminotypes in heat and control treatments. Seventeen unique aminotypes were detected in coral fragments exposed to heat, compared to 1 unique aminotype in control fragments. (b, h) Diversity (Shannon’s diversity index, H) of aminotypes increased in fragments exposed to heat compared to control fragments, and over time (d, h); variation also appeared to occur among colonies (f). (c, i) Estimated aminotype richness increased in fragments exposed to heat, but not over time (e, i). (g) Mean aminotype richness also appeared to vary among colonies. Linear mixed effects model (LMM) results for Shannon’s diversity index (h) and estimated richness (i). Shannon’s diversity index values were based on sequencing data rarefied to 59 837 amino acid sequences per sample; richness was estimated by repeated random subsampling of unrarefied data. Coral-symbiont combinations were pooled for these analyses. Colonies 1, 4 and 5 are *Pocillopora verrucosa* (containing *Cladocopium pacificum*), colonies 2 and 3 are *P. ligulata* (containing *C. latusorum*).

Alpha diversity (Shannon’s index, H) of aminotypes was positively associated with the heat treatment (F=4.87, p=0.03) and time (F=4.91, p=0.03) in our linear mixed effects model (LMM; Figure 3b, d, h). There was no significant interaction between heat and time (F=0.57, p=0.46), nor was there a significant difference between coral-symbiont species (F=0.44, p=0.55). On average (± SE), Shannon’s index was 27% higher in heat-treated fragments (1.1 ± 0.6) than in control fragments (0.8 ± 0.6; Figure 3b). Within individual timepoints, heated fragments had 4-93% higher mean H values than controls (0.9-1.3 versus 0.7-1.0, respectively; Figure 3d), but individual comparisons were not significant. Means of Shannon’s index per colony ranged between 0.5-1.8 for heat-treated fragments and 0.1-1.6 for control fragments (Figure 3f).

There was a significant positive association between viral aminotype richness and heat treatment (F=5.70, p=0.02), but no effect of time (F=0.35, p=0.56) nor coral-symbiont species (F=0.001, p=0.99) in our LMM (Figure 3c, e, i). There was no significant interaction between treatment and time (F=0.10, p=0.75). On average, aminotype richness was 16% higher in heat-treated (37.2 ± 2.0) than control fragments (32.0 ± 2.1; Figure 3c). At individual timepoints, heat-treated fragments had 7-23% higher mean aminotype richness than controls (34.8-39.1 versus 29.2-35.4, respectively; Figure 3e), but these differences were not significant. Mean aminotype richness for individual colonies (using timepoints as replicates) ranged between 28.0-44.9 for heat-treated fragments and 19.5-44.6 for control fragments (Figure 3g).

### Twenty-two aminotypes had higher relative abundances in heat-treated fragments

DESeq2 analysis revealed 28 aminotypes had significantly altered relative abundances in heat-treated or control fragments (Figure 4). Twenty-two aminotypes had higher relative abundances in heated fragments, whereas 6 aminotypes had higher relative abundances in controls. Many (16/28) of the differentially abundant aminotypes were identified in both coral-symbiont species (aminotype names without a circle or asterisk, Figure 4). Ten aminotypes were differentially abundant at multiple (2-4 out of 5) timepoints throughout the experiment; all ten of these were identified in both coral-symbiont species. Aminotypes 29, 38 and 98 were significantly more abundant in the heat treatment in 4 out of 5 sampled timepoints (from 12 to 108 h); aminotype 93 was more abundant in the final 3 timepoints (from 24 to 108 h).

**Figure 4.**
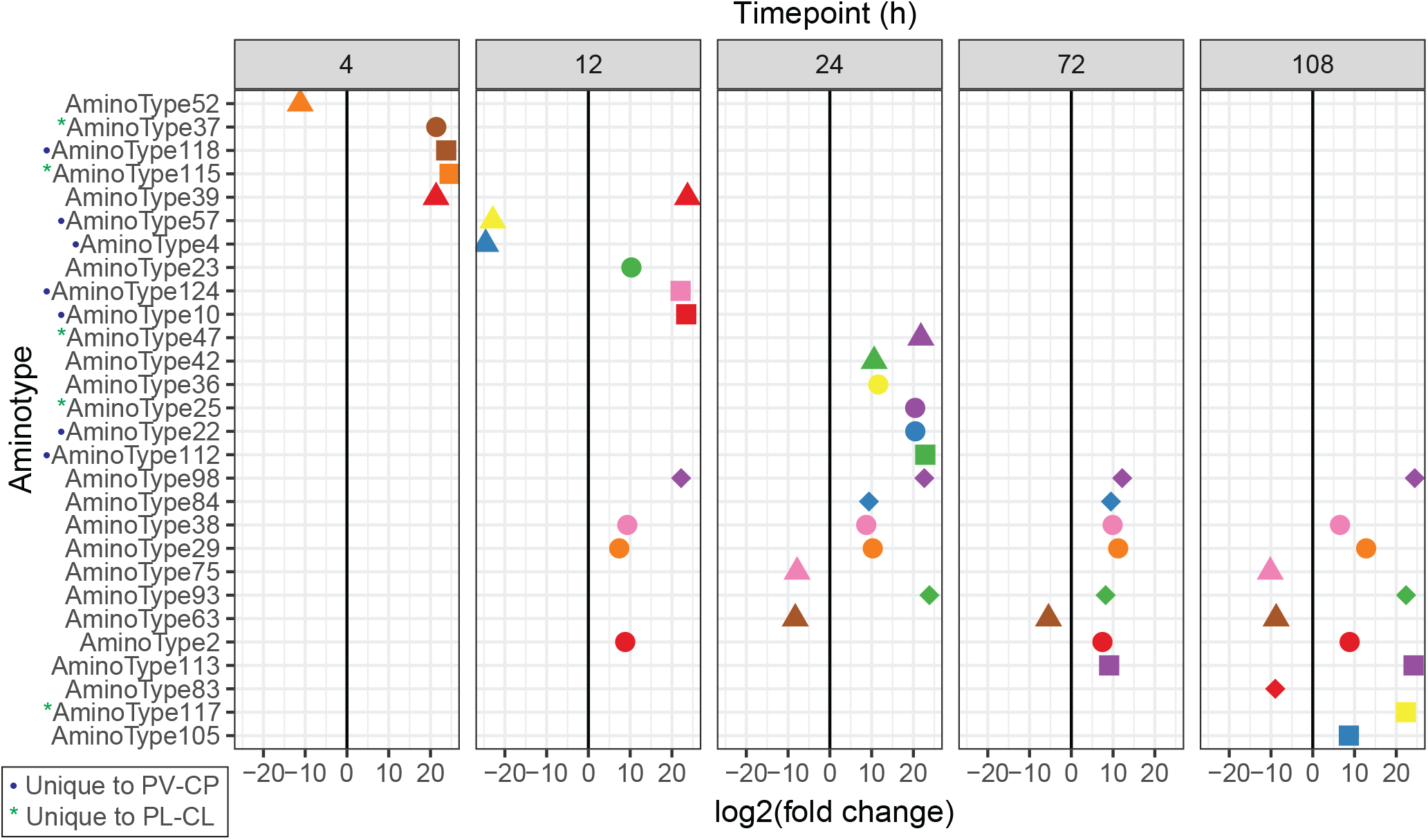
Differentially abundant Symbiodiniaceae-infecting dinoRNAV major capsid protein (*mcp*) aminotypes (unique amino acid sequences) across the experiment. Twenty-eight viral aminotypes (each represented as a point with a unique shape and color) were differentially abundant across control and heated fragments of the coral-symbiont species *Pocillopora verrucosa*-*Cladocopium pacificum* (PC-CP) and *P. ligulata*-*Cladocopium latusorum* (PL-CL). Of these, 22 aminotypes occurred at higher relative abundances in heat-treated fragments (right of the 0 lines) and 6 had higher relative abundances in control fragments (left of the 0 lines). Ten aminotypes were differentially abundant at multiple (2-4) timepoints throughout the experiment. This analysis was conducted using DeSeq2 on a non-rarefied dataset including both coral-symbiont species. Blue circles indicate aminotypes unique to PV-CP; green asterisks indicate aminotypes unique to PL-CL. DESeq2 analyses were run separately for each time point (t_(h)_= 4, 12, 24, 72, 108, see Methods for more details).

### Dispersion of dinoRNAV *mcp* amino acid sequences increased in heat-treated fragments

Dispersion (measured as distance to centroid) of dinoRNAVs was positively associated with heat treatment (Figure 5a, b, e; F=9.28, p=0.004) in the LMM. There was no effect of time (F=0.23, p=0.95) nor coral-symbiont species (F=0.20, p=0.68), and no interaction between treatment and time (F=0.48, p=0.75). Overall, mean (±SE) dispersion was 62% higher in the heat treatment when all colonies and timepoints were pooled (heat: 0.23 ± 0.13; control: 0.14 ± 0.09). A trend of increasing dispersion of dinoRNAVs (32-100% higher) was observed in heat-treated samples at individual timepoints (Figure 5c; heat: 0.2-0.29; controls: 0.12-0.17), but individual comparisons were not significant. Mean dispersion of individual colonies (across timepoints) was 11-93% higher per colony in heat-treated fragments (ranging from 0.13-0.37 in heat-treated fragments versus 0.1-0.24 in controls), but individual comparisons were not significant (Figure 5d).

**Figure 5.**
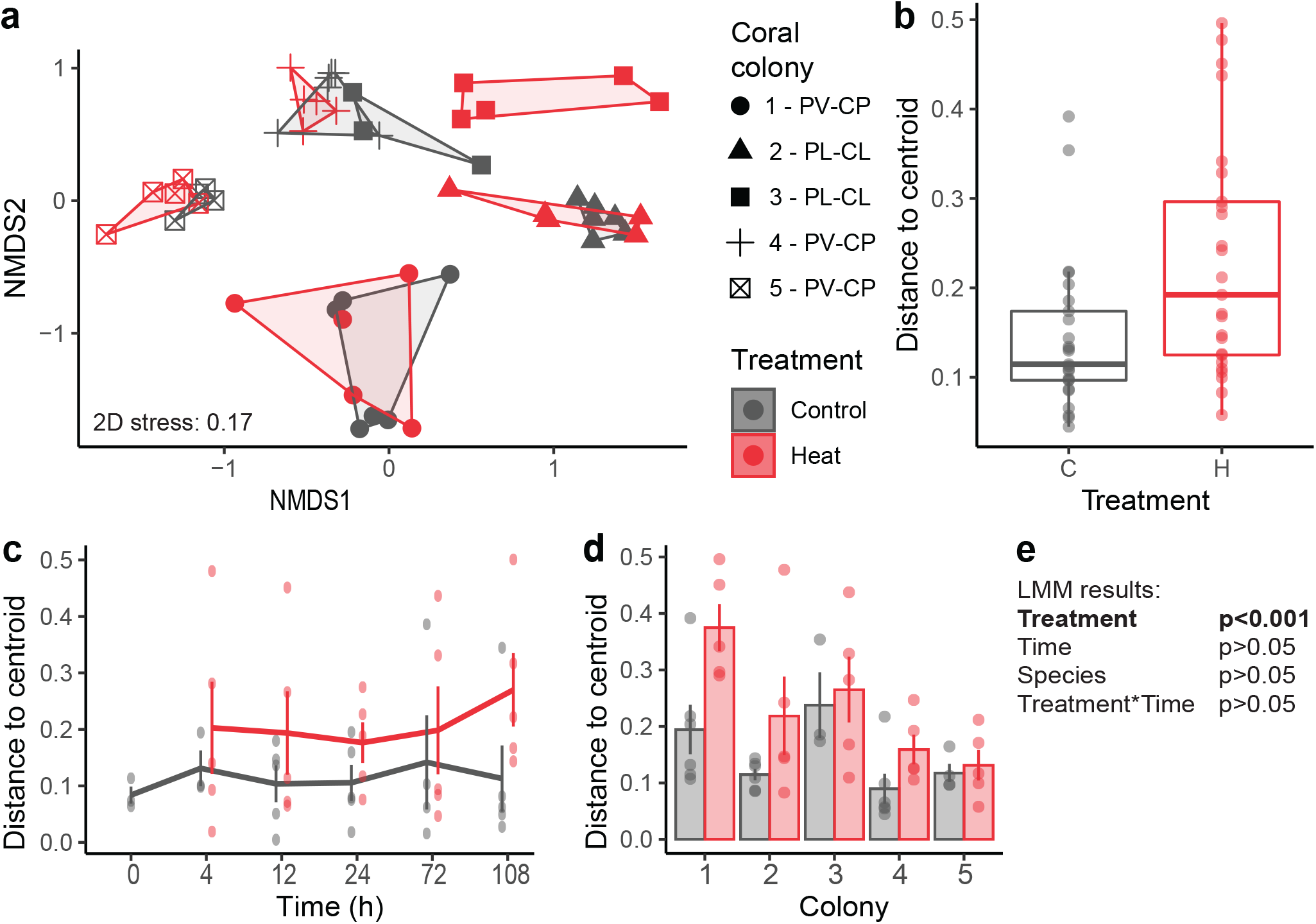
Dispersion of Symbiodiniaceae-infecting dinoRNAV major capsid protein (*mcp*) gene aminotypes (unique amino acid sequences) associated with coral fragments in heat-treated (H) versus control (C) conditions. (a) A non-metric Multidimensional Scaling (nMDS) plot depicts that dinoRNAVs differ by coral colony ID, coral-symbiont species (*Pocillopora verrucosa-Cladocopium pacificum*, PV-CP; *P. ligulata-C. latusorum*, PL-CL), and treatment. (b) Dispersion of dinoRNAV aminotypes was higher in heat-treated fragments. (c) Mean (±SE) aminotype dispersion was consistently higher over time in heat-treated fragments and (d) varied among coral colonies. (e) Results of a linear mixed effects model testing the effect of treatment and time on dispersion. Centroids were calculated separately for each colony in control and heat-treated conditions based on Bray-Curtis distances from square-root-transformed rarefied data. Dispersion was quantified by measuring the distance of each sample to its centroid.

### DinoRNAV *mcp* and *RdRp* transcripts were detected in all metatranscriptomes

RNA sequencing resulted in 436 532 reads with similarity to reference viral sequences in the RVDB. A mean of 1.17% ± 1.04 (SD) of viral reads showed homology to the putative RNA-dependent RNA polymerase (*RdRp*) of dinoRNAVs previously reported in association with *Cladocopium* C1 (Table S2; [27]). Overall, best hits to the *RdRp* of *Cladocopium* C1-infecting dinoRNAVs were identified in all analyzed metatranscriptomes (colonies 1, 4, 5; Table S3). These hits were the tenth-most common viral hits overall, and the fourth most common putative eukaryotic virus hits (Table S2). Other dinoRNAV-like reads matched best to two *mcp* sequences of *Cladocopium* C1-infecting dinoRNAV [27], comprising 0.06% ± 0.05 (AOG17586.1) and 0.04% ± 0.03 (AOS87317.1) of the viral transcripts, respectively (Table S3).

RnaSPAdes produced 214 309 contigs, of which 11 804 showed homology to virus reference protein sequences from the RVDB with DIAMOND BLASTx. A total of 128 contigs (lengths 211 - 2 631 nucleotides) showed homology to reference dinoRNAV ORFs from Levin et al. [27]. Amino acid-based alignment of the 38 recovered *mcp* contigs (and amplicons) generated in this study to the dinoRNAV *mcp* ORF from Levin et al. [27] revealed that although the dinoRNAV *mcp* gene is variable, it also contains five relatively conserved regions (CR I-V in Figure S3). No significant insertions or deletions were apparent in the *mcp* contigs or amplicon sequences recovered in this study (Figure S3).

## Discussion

Viruses can have diverse impacts on hosts, ranging from antagonistic to beneficial [24, 68–70]. Efforts to understand how viral infections impact coral colonies have been stymied by the lack of (1) a high-throughput approach to track a viral lineage in colonies across an acute stress event; and established cultures of viruses associated with corals and their symbionts (for use in viral addition experiments). This study tracks a group of Symbiodiniaceae-infecting viruses, the dinoRNAVs, in a controlled experiment to interrogate how infection responds to temperatures associated with bleaching. DinoRNAVs were detected from all five colonies examined, and from every heat-treated coral fragment, but only some controls. These observations, and detections of higher alpha diversity of dinoRNAV *mcp* aminotypes in heat-treated fragments, along with increased aminotype dispersion, unique aminotypes, and greater relative abundances of specific aminotypes, together strongly indicate that dinoRNAV infections were more active in heat-treated fragments. DinoRNAV responses were detectable within a single day, more quickly than the several weeks over which bleaching signs typically manifest. If viral infection of Symbiodiniaceae cells is enhanced (e.g., increased production, accumulation of viral diversity) during thermal anomalies *in situ*, then cumulative viral activity, if maintained over weeks, may disrupt coral-symbiont partnerships and potentially modulate some bleaching responses on reefs.

### DinoRNAVs as a common, persistent virus of Symbiodiniaceae

DinoRNAV *mcp* genes were detected in all experimental colonies (N=5), and in most to all fragments per colony (73-100% of fragments, N=8/11 – 11/11 fragments per colony) using the gene amplicon sequencing method. DinoRNAV genes have additionally been reported from colonies of seven other stony coral species across the Atlantic and Pacific Oceans (Table 1, Figure 1), suggesting that these viruses are commonly associated with coral microbiota. In this experiment, viral aminotype compositions (and potential viral quasispecies) differed among coral-symbiont species and colonies, but were similar within the fragments of a given colony, despite maintenance in separate aquaria (Figure 2). These results suggest that, under ambient conditions, dinoRNAV populations are driven strongly by within-species and within-colony factors. RNA viruses rely on RNA-dependent RNA polymerase (*RdRp*) for replication, which is relatively error-prone. Therefore, individual viral progenitors infecting Symbiodiniaceae cells in a given colony may each produce a variety of genetically distinct viral “progeny” during a single replication cycle [35, 71]; hence, the observed pattern of species- and colony-specificity in dinoRNAV aminotypes likely results from diversification within ‘quasispecies’ and (potentially) subsequent purifying selection [35, 72–74]. This production of a ‘mutant cloud’ of dinoRNAV diversity may even help ensure these viruses are successful (at the population level) at infecting Symbiodiniaceae under changing environmental conditions [37]. As a next step, future works can quantify the extent to which dinoRNAV diversity is homogeneously distributed across entire coral-symbiont colonies at a given timepoint using multiple markers (e.g., *mcp* and *RdRp*).

Healthy corals contain millions of Symbiodiniaceae cells per cm^2^ of coral tissue; our results suggest that Symbiodiniaceae *in hospite* may be infected by one or perhaps several dinoRNAV quasispecies (e.g., aminotypes 30 and 52 in colony 2, Figures 2 and S2) at any given time. Recent surveys of free-living marine microbial communities reported that viral infections may occur in ∼33% [75, 76] to over 60% [77] of marine microorganisms, and many individual cells may be infected by multiple distinct viruses at a given time [75, 78]. Given this, the detection of multiple dinoRNAV *mcp* genes or quasi-species from a single Symbiodiniaceae cell *in hospite* might be expected; single cell RNA-Seq should be used to test this possibility.

The dinoRNAV *mcp* gene was not amplifiable from DNA extractions via PCR [79–81], and *mcp* transcripts from the metatranscriptomes did not contain significant insertions or deletions (compared to the full *mcp* gene assembly [27], figure S3). This indicates that the dinoRNAV *mcp* sequences reported here constitute exogenous viral infections, rather than endogenous viral elements (EVEs; [62]) within their host’s genome. Considering that the gene amplicons reported here were generated from unfractionated coral tissue, dinoRNAV *mcp* detections in this study could potentially arise from four sources: RNA genomes within intact dinoRNAV capsids (*sensu* [29]); *mcp* genes that are being expressed in host cells during a dinoRNAV replication cycle; free dinoRNAV genomes that may occur as extrachromosomal RNA in a latent [26] or carrier state similar to pseudolysogeny [82, 83]; and /or chimeric RNA viruses with analogous *mcp* gene sequences [61]. However, the identification of highly abundant transcripts with homology to Symbiodiniaceae-associated dinoRNAV *RdRp* genes in the metatranscriptomes (Table S2, S3) indicates that *mcp* gene amplicons in this study are most parsimoniously interpreted as derived from dinoRNAVs [79, 80].

The sequencing approaches employed here limit our ability to discern amongst the potential sources of viral *mcp* gene detections, as does the dearth of information available on dinoRNAV replication cycles in coral symbionts. Approaches such as single cell RNA-seq (e.g., [84]) of Symbiodiniaceae cells, as well as sequencing methods designed to identify and characterize infective viruses (similar to viral tagging [85], adsorption sequencing [86]) represent critical next steps in assessing the dynamics of dinoRNAV infections within individual Symbiodiniaceae cells.

### Switching from persistent to more productive infection under stress

Many viruses switch between strategies upon environmental changes [24, 87–91]. RNA virus infections can range from exclusively lytic [30, 94, 95], to infections that are ‘persistent’ or ‘chronic’ and do not immediately kill the host [96, 97]. Persistent RNA viruses of plants, for example, may show altered activity based on seasonality [98, 99], temperature [99, 100], or other environmental factors [101, 102]. Similarly, marine viruses can switch between latent and productive cycles in response to host-related or environmental triggers [8, 92, 93]. For persistent viruses of Symbiodiniaceae, a variety of mechanisms may modulate viral infection strategies; the role of temperature in triggering persistent infections to become more productive has received particular attention due to the link between temperature and coral bleaching [8, 11, 25, 27, 93,101].

We identified 22 aminotypes that were present at higher relative abundances in heat-treated coral fragments (Figure 4), and 17 aminotypes were unique to these fragments (Figure 3a, c). We interpret that these findings indicate a switch from persistent to more productive infections by some dinoRNAV strains (or quasispecies). Of particular interest, aminotypes 29, 38, 93 and 98 exhibited sustained increases in relative abundance in heat-treated corals starting at 12-24 hours (Figure 4). Screening for these dinoRNAV *mcp* aminotypes in additional Symbiodiniaceae populations (and sequencing libraries) is of interest.

The observation of a rapid increase in dinoRNAV aminotype diversity in some colonies during the onset of thermal stress is consistent with a previous report of increased (but overall low) abundance of dinoRNAV transcripts in the Caribbean coral *Montastrea cavernosa* following exposure to elevated temperatures for 12 hours [25]. Further, a thermosensitive culture of *Cladocopium* C1 exhibited high expression of transcripts with best hits to dinoRNAV genes under ambient temperatures, while a thermotolerant *Cladocopium* C1 culture did not, suggesting that dinoRNAVs may modulate resistance to bleaching in some species or populations of Symbiodiniaceae [27]. Similarly, viral metagenomes from bleached pocilloporid colonies *in situ* contained significantly more eukaryotic virus sequences than unbleached, apparently healthy colonies [11]. Experiments with the cricket paralysis virus—a +ssRNAV with similar genome architecture to dinoRNAVs in Symbiodiniaceae cultures—also showed increased viral replication after two hours when temperatures were raised by 5°C [102].

### Colony-specific dinoRNAV responses to elevated temperatures

Since higher temperatures increase host cell enzymatic activity, increased viral production and accumulation of mutations within quasispecies exposed to heat stress might be expected based on thermodynamics alone [104]. However, dinoRNAV responses to elevated water temperatures differed among individual coral colonies (Figures 2b, 5a), even in two colonies (3 and 4) that were both dominated by aminotype 1 in control aquaria. While dinoRNAVs in heat-treated fragments from colonies 1 and 3 exhibited strong shifts in putative quasispecies compositions, such changes were less pronounced in the other colonies. These findings suggest that dinoRNAV strains (or quasispecies) in colonies 1 and 3 were more responsive to heat stress. Coral colonies generally exhibit heterogeneous resistance to coral bleaching (e.g., [11, 105, 106]), and bleaching susceptibility in pocilloporids has been correlated to differential communities of eukaryotic viruses [11]. Subsequent experiments that extend multiple weeks to sample both the onset of thermal stress and the onset of bleaching signs are a critical next step in understanding how dinoRNAV dynamics relate to colony health trajectories and relative bleaching resistance.

## Conclusions

This is the first study to characterize the dynamics of Symbiodiniaceae-infecting dinoRNAVs in coral colonies exposed to ecologically relevant bleaching temperatures. We identified dinoRNAVs in each sampled coral; temperature stress elicited rapid changes to dinoRNAV diversity and composition, and a subset of viral aminotypes were significantly associated with heat-treated fragments. Multiple lines of evidence suggest that dinoRNAVs are common in pocilloporid corals as persistent, exogenous infections of Symbiodiniaceae. Environmental stress may increase the productivity of these viruses, potentially impacting colony symbiotic status (if stress is prolonged). Overall, these findings add to the growing body of literature demonstrating that viruses of microorganisms affect emergent phenotypes of animal and plant holobionts, and may modulate holobiont responses to changing environmental conditions.

## Acknowledgements

The authors express their sincere appreciation to Kira Turnham and Dr. Todd LaJeunesse for resolving coral and Symbiodiniaceae species in this study, and to Drs. Rebecca L. Vega Thurber, Andrew R. Thurber, and Craig E. Nelson for discussions and logistical support related to the experiment. Many thanks to Rebecca L. Maher and J. Grace Klinges for help with sampling, and to Dennis Conetta for assistance with DNA and RNA extractions. We additionally thank Mark Dasenko at Oregon State University’s Center for Genome Research & Biocomputing (Corvallis, OR) for his support in designing the sequencing methods. Lastly, we also thank two anonymous reviewers for their suggestions and comments that improved the manuscript. Financial support was provided by a Sigma-Xi Grant-in-aid of Research to C.G., a U.S. National Science Foundation award (OCE #1635798) to A.M.S.C. and an Early-Career Research Fellowship (#2000009651) from the Gulf Research Program of the National Academies of Sciences to A.M.S.C.

## Author contributions

C.G., L.H.K. and A.C. conceived of the experiment; C.G., L.H.K., and A.J.V. developed the methods with support from A.C.; C.G., L.H.K., R.B., and A.C. conducted the experiments and processed samples; C.G. led data analysis, with contributions by all authors; C.G. wrote the first draft of the manuscript, with contributions by all authors.

