## Supplementary Material for "Thermal stress triggers productive viral infection of a key coral reef symbiont"

### Supplementary Methods

##### **Experimental design**

Five *Pocillopora-Cladocopium* (coral-symbiont) colonies were collected in the shallow backreef (1-2 meter depth) of Mo’orea, French Polynesia (17º 32' S 149º 50' W) during the dry season in September 2017 (Figure S1). The total number of colonies used in the experiment was limited to five because that was the maximum number of colonies that could be maintained in the experimental aquariums while maintaining plenty of separation between coral fragments. Collected colonies were placed in flow-through water tables, fragmented, and allowed to acclimate for 24 hours. All fragments were then distributed randomly across eight experimental 75 L aquarium tanks, four of which were randomly designated as control tanks, and the other four were designated as experimental ‘heat’ tanks. Fragments were assigned to control or heat tanks so that six (control treatment) or five (heat treatment) fragments from each colony were distributed across control or heat treatments tanks respectively. Treatment assignment was conducted using a blinded design so that the handler did not know which treatment each fragment would be assigned to. All tanks were shaded from sunlight, provided with seawater passed through a coarse sand filter at a rate of 90 L h^-1^, and temperature and light intensities were continuously recorded using data loggers (ONSET, Bourne). To mimic local peak temperatures associated with coral bleaching, four randomly selected tanks were each equipped with two 300W water heaters (Finnex, Chicago). The average temperature throughout the experiment was 28.2˚C in control aquaria (similar to ambient reef water temperatures) and 30.3˚C in heat-treated aquariums Thus, there was an average difference of 2.1˚C between treatments (Figure S1).

##### **Coral fragment colorimetric analysis**

To characterize the effect of the heat treatment on Symbiodiniaceae cell densities (a metric of coral health and bleaching status), we measured color values for each coral fragment as a proxy for Symbiodiniaceae chlorophyll concentrations [1, 2]. The color balance was manually adjusted to be equivalent across all images in Adobe Photoshop (v. 21.1.3), and the hue, saturation and brightness values of each fragment were measured in an area of 101x101 pixels [1]. The color values for each fragment at the time of sacrifice were then divided by the values at the start of the experiment, resulting in values describing the change in color in each fragment from the start of the experiment to the time of sacrifice.

##### **Sampling and DNA and RNA extractions**

Samples (which included coral animal tissue, Symbiodiniaceae cells and viruses) were collected from each coral fragment using sterilized bone clippers and transferred to 15 ml conical tubes with 5 ml DNA/RNA shield (Zymo Research, Irvine, CA, USA), Lysing Matrix A (garnet) and 0.6 cm ceramic beads (MPBio, California). All tubes were vortexed for 20 minutes at maximum speed to disrupt tissues and stored at -20 ºC.

DNA and RNA were extracted from vortexed tissue slurry using the ZymoBIOMICS DNA/RNA Miniprep Kit according to the manufacturer’s instructions but with an additional enzyme digestion step to improve viral RNA yields. Briefly, 30 µl lysozyme (10 mg ml^-1^), and 1.8 µl of each lysostaphin (4 KU ml^-1^) and mutanolysine (50 KU ml^-1^) were added to 300 µl of coral tissue in DNA/RNA shield and incubated at 37 ºC for one hour. Then, 30 µl of proteinase K digestion buffer with 15 µl proteinase K (20 mg ml^-1^) was added and incubated at 50 °C for one hour. DNA and RNA were then extracted in parallel according to the manufacturer’s instructions and a 15 min DNase step was included to remove potential DNA contamination in isolated RNA; the extracted DNA and RNA was then eluted in 100 µl sterile water. 2.5 µl of RiboGuard RNAse inhibitor (Lucigen, Middleton, WI, USA) was added to extracted RNA to prevent degradation. The concentration and quality of the DNA and RNA was assessed using a NanoDrop spectrophotometer (Thermo Fisher Scientific, Waltham, MA, USA) before storage at -80 °C. DNA concentrations ranged from 40 to 90 ng µl^-1^. RNA concentrations ranged from 20-100 ng µl^-1^.

##### **DinoRNAV major capsid protein (*mcp*) gene amplicon sequencing**

Complementary DNA (cDNA) was synthesized from 5 µl of extracted RNA (100-4500 ng) with SuperScript III Reverse Transcriptase (Thermo Fisher Scientific) following the manufacturer’s instructions. To prime the template RNA, 120 ng of Random Primers (Thermo Fisher Scientific) was added, and RNaseOUT (Thermo Fisher Scientific) was used to prevent RNA degradation during synthesis. The dinoRNAV major capsid protein (*mcp*) gene was amplified from cDNA using a nested PCR protocol with degenerate primers [3]. For the first round of PCR we used 10 µM of each of the primers HcUniv-01F [5`TCCTTGTWTRYWKGATGCKTTTCA`3] and HcUniv-01R [5`MGCCAARTCASWCATATTAAAWGGCA`3]. We used the Qiagen Multiplex Kit (Qiagen, Germantown, MD, USA) in 20 µl reactions with the following thermocycler program: 95°C for 15min, [94°C for 30 s, 60°C for 90 s, 72°C for 90 s] 30 cycles, 72°C for 10 min. For the second round of PCR, we used 10 µM of each of the primers HcUniv-02F  [5'TCGTCGGCAGCGTCAGATGTGTATAAGAGACAGYTKCCTCGASCTRYTGGWCC'3] and HcUniv-01R [5'GTCTCGTGGGCTCGGAGATGTGTATAAGAGACAGMGCCAARTCASWCATATTAAAWGGCA'3] with Illumina adapter overhangs (underlined above). The 2x KAPA HiFi HotStart ReadyMix (Roche, Switzerland) was used with 1 µl of product from PCR 1 as template in a 25 µl reaction volume. The thermocycler program was the same as for the first round of PCR, but with 25 cycles instead of 30.

The PCR product was cleaned using Mag-Bind TotalPure NGS (Omega Bio-tek, Norcross, GA) and indexed using the Nextera XT Index kit (illumina, San Diego, CA). Samples were then cleaned again and quantified with a BioTek Synergy H1 Microplate Reader with the Quant-iT 1x dsDNA HS kit (Invitrogen, Carlsbad, CA), and subsequently normalized and pooled. Amplicon sizes were checked using the TapeStation 4200 with a HS-D5000 tape (Agilent, Santa Clara, CA) and concentrations were measured with qPCR using a KAPA biosystems library quantification kit (Roche, Switzerland).

##### **Processing of dinoRNAV *mcp* gene reads**

Initial quality checks of raw read libraries were done using the program fastQC v0.11.9. Next, adapter sequence detection and removal, read length filtering and read quality filtering were conducted using the software fastp v0.20.1 [4] with default parameters. From the remaining reads, primer sequences were trimmed using the program bbduk.sh from the BBTools bioinformatics suite (v37.88; sourceforge.net/projects/bbmap/). The cleaned paired-end reads were then merged using the program vsearch (v2.14.2; [5]) and any merged reads with lengths <400 nucleotides were removed with fastp v0.20.1. All merged reads with a length of 422 nucleotides were extracted and dereplicated into unique sequences using vsearch v2.14.2. With the unique sequences, amplicon sequence variant (ASV) generation and subsequent *de novo* chimera detection and removal were performed using vsearch with the UNOISE3 and UCHIME3 algorithms, respectively [6, 7]. ASVs containing stop codons were removed (11 of 273 total ASVs) and the remaining ASVs were translated into amino acid sequences. Retained translated ASVs were clustered by 100% amino acid identity into “aminotypes” (unique amino acid sequences) with the program CD-HIT v4.8.1 [8]. Merged reads with lengths between 400 and 422 base pairs (from the total dataset) were translated and aligned to the aminotypes using DIAMOND BLASTx v2.0.11 [9] to produce a counts table.

**Metatranscriptome sequencing and bioinformatics processing**

Metatranscriptomes were generated from five samples of *Pocillopora verrucosa* harboring *Cladocopium pacificum* (t_(h)_=0 control for colony 1, t_(h)_=3 heat for colony 4, t_(h)_=5 control and heat for colony 5, t_(h)_=4 control for colony 5). Ribosomal RNA was removed from RNA libraries using equal parts plant leaf, human/mouse and bacteria Ribo-Zero kits (Illumina). cDNA was then synthesized and sequenced on a HiSeq 3000 instrument using 150 bp PE chemistry at Oregon State University.

Adapters and low-quality reads were removed from the generated RNA sequence libraries using fastp v0.20.1 [4]. For initial assessment of dinoRNAV-derived reads within sequenced samples, cleaned reads were merged with bbmerge.sh and aligned to the proteic version of the Reference Virus Database (v21) using DIAMOND BLASTx v2.0.11 [9]. The relative abundances of hits to each viral gene were then calculated (see Table S2 and S3 for results).

Cleaned reads were then separated into forward and reverse reads, normalized using bbnorm v38.79 [10], and assembled into contigs using rnaSPAdes v3.13.0 [11]. Contig taxon identity and function were inferred by DIAMOND BLASTx (v 2.0.11) alignment (minimum bitscore=50; minimum ORF length=30 aa; minimum percent identity= 30) to a hybrid database containing the uniprot database and the Reference Virus Database [12].

The program Prodigal (v.2.6.3) was then used to predict open reading frames and protein sequences from all dinoRNAV-like transcripts, and protein annotation was conducted via another round of DIAMOND BLASTp alignment to the hybrid database. Confirmed dinoRNAV major capsid protein (*mcp*) amino acid sequences from the identified dinoRNAV transcripts were then aligned with the dinoRNAV *mcp* sequence from Levin et al. [13] using the program muscle v.5 [14].

#### Supplementary Tables and Figures

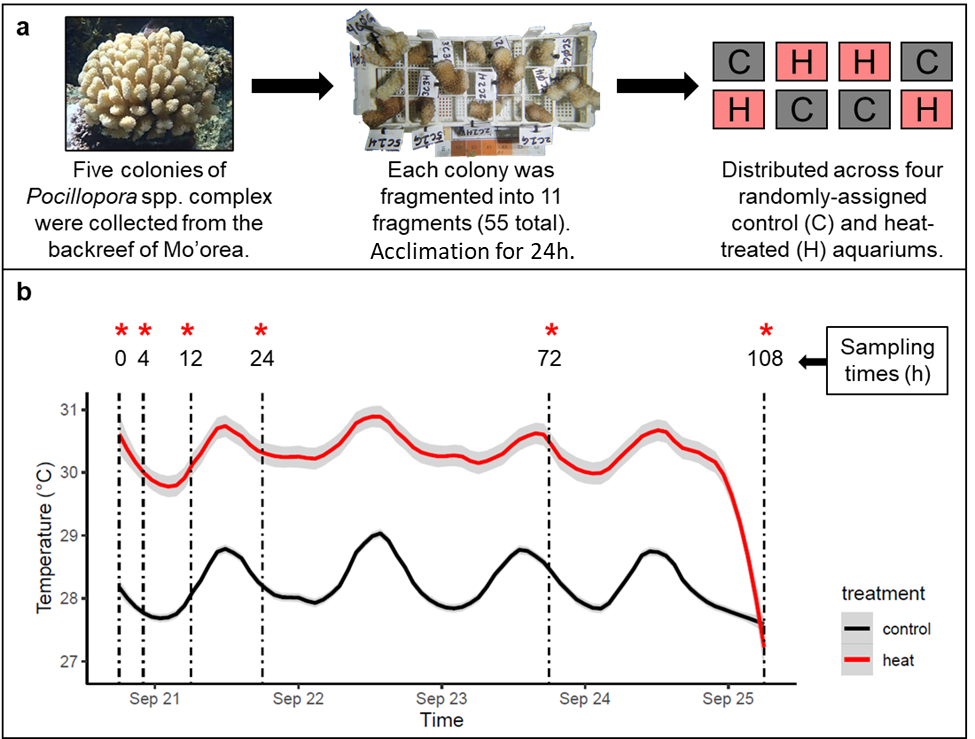
**Figure S1**	Experimental design: five *Pocillopora-Cladocopium* (coral-symbiont) colonies [15] were collected in the shallow backreef (1-2 meter depth) of Mo’orea, French Polynesia during the dry season in September 2017. Colonies were placed in flow-through water tables, fragmented into 11 fragments each, and allowed to acclimate for 24 hours. All fragments were then randomly distributed across eight 75 L aquarium tanks, half of which contained water heaters (a). The overall average temperature was 28.2˚C in control aquariums (similar to ambient reef water temperatures) and 30.3˚C in heated aquariums, resulting in an average temperature difference of 2.1 ˚C between treatments (b). All fragments were photographed at each timepoint (0, 4, 12, 24, 72, 108 h) and one fragment from each colony per treatment was sacrificed for sequencing of the dinoRNAV major capsid protein (*mcp*) gene (red asterisks).

**Table S1.**	Overview of dinoRNAV major capsid protein (*mcp*) aminotype (unique amino acid sequences) alignment to a custom database using Diamond blast. The custom database included all sequences from the reference viral database (RVDB; [12]) and the aminotypes generated by reprocessing dinoRNAV *mcp* nucleotide data generated from Great Barrier Reef corals in [3]. All 124 aminotypes in this study aligned to one of 13 reference amino acid sequences in [3]. ‘N_unique aminotypes (this study)’ lists the number of unique amino acid sequences in our dataset with a best hit to a given ‘Reference sequence’ in our hybrid database. ‘Mean ID (%)’ indicates the mean percentage of identical characters among sequences from this study and the reference sequence. ‘Mean bitscore’ and ‘Mean e-value’ indicate mean bitscores and e-values of the alignments, respectively. Mean ID (%) ranged from 52.1-99.3, mean bitscores ranged from 152.5-276.9; and mean e-values ranged from 9.6·10^-73^-2.7·10^-35^.

| *N_unique aminotypes*  (this study) | *Reference sequence* | *Mean ID (%)* | *Mean bitscore* | *Mean e-value* |
| --- | --- | --- | --- | --- |
| 4 | Pdamicornis_AminoType4 | 98.40 | 275.13 | 1.43E-71 |
| 8 | Pdamicornis_AminoType1 | 89.98 | 258.41 | 1.36E-66 |
| 3 | Pdamicornis_AminoType3 | 87.33 | 253.70 | 2.53E-62 |
| 11 | Plutea_AminoType5 | 79.24 | 223.65 | 2.43E-41 |
| 1 | Atenuis_AminoType3 | 77.70 | 226.50 | 1.50E-57 |
| 3 | Pcylindrica_AminoType1 | 75.10 | 220.33 | 4.33E-51 |
| 3 | Plutea_AminoType10 | 70.23 | 202.47 | 4.53E-50 |
| 2 | Plutea_AminoType17 | 66.30 | 197.40 | 1.27E-48 |
| 3 | Plutea_AminoType4 | 65.47 | 200.03 | 1.68E-49 |
| 5 | Gfascicularis_AminoType1 | 60.52 | 169.94 | 1.00E-36 |
| 21 | Plutea_AminoType13 | 55.20 | 156.99 | 2.18E-36 |
| 56 | Plutea_AminoType21 | 55.13 | 157.45 | 2.38E-36 |
| 4 | Plutea_AminoType11 | 53.40 | 154.85 | 6.38E-36 |

**Table S2.**	Top 20 most abundant viral read alignments from five metatranscriptomes generated from three colonies of *Pocillopora verrucosa-Cladocopium pacificum* to amino acid sequences in the reference viral database (RVDB; [12]). Hits to genes of putative eukaryotic viruses are **bolded**. Other hits are to genes of phages or unclassified viruses. Hits to the genome assembly of Symbiodiniaceae-infecting dinoRNAVs are *italicized* [13]. All metatranscriptomes were normalized before calculating mean read abundances. SD: Standard deviation.

| *Accession #* | *Biological entity* | *Gene* | *Mean read abundance (%)* | *SD* |
| --- | --- | --- | --- | --- |
| YP_009126958.1 | Eel River basin pequenovirus | Putative major capsid protein | 22.82 | 10.94 |
| YP_009126954.1 | Eel River basin pequenovirus | Putative replication initiation protein | 9.58 | 4.59 |
| **CCQ19277.1** | **Cotesia sesamiae Kitale bracovirus** | **Histone H4-like** | **6.78** | **3.44** |
| **ABH10013.1** | **Cotesia glomerata bracovirus** | **Putative histone 4** | **6.60** | **2.65** |
| YP_009126956.1 | Eel River basin pequenovirus | Putative minor capsid protein | 5.71 | 2.67 |
| **YP_009465730.1** | **Dishui lake phycodnavirus 1** | **Hypothetical protein** | **3.64** | **1.91** |
| ACF23597.1 | Uncultured marine virus | Photosystem II protein D1 | 1.82 | 0.71 |
| ABF59588.1 | Uncultured virus | Photosystem D1 protein | 1.66 | 0.77 |
| AAU84538.1 | Uncultured marine virus | Photosystem II D1 protein | 1.28 | 0.54 |
| ***AOG17585.1*** | ***Symbiodinium +ssRNA virus TR74740 c13_g1_i1*** | ***putative RNA-dependent RNA polymerase*** | **1.17** | **1.04** |
| ACF23951.1 | Uncultured marine virus | Photosystem II protein D1 | 0.92 | 0.23 |
| QHT80240.1 | Viral metagenome | Hypothetical protein | 0.77 | 0.27 |
| **AEE09495.1** | **Cotesia vestalis bracovirus** | **Histone** | **0.61** | **0.32** |
| QHT05404.1 | Viral metagenome | Hypothetical protein | 0.61 | 0.23 |
| **CAA42007.1** | **Epstein-Barr virus** | **Epstein-Barr virus small RNA associated protein** | **0.59** | **0.27** |
| AAU84539.1 | Uncultured marine virus | Photosystem II D1 protein | 0.54 | 0.13 |
| **YP_184795.1** | **Cotesia congregata bracovirus** | **Hypothetical protein** | **0.47** | **0.23** |
| ACF24065.1 | Uncultured marine virus | Photosystem II protein D1 | 0.43 | 0.10 |
| QHT36437.1 | Viral metagenome | Hypothetical protein | 0.39 | 0.12 |
| QHS93229.1 | viral metagenome | Hypothetical protein | 0.35 | 0.14 |

**Table S3.**	Relative abundances (A) and alignment quality values (B) of viral reads in each metatranscriptome with hits to three reference dinoRNAV amino acid sequences (AOG17585.1: Putative RNA-dependent RNA polymerase, *RdRp*; AOG17586.1 and AOS87317.1: Putative major capsid protein, *mcp*) from Levin et al. [13]. Metatranscriptomes were generated from fragments of three coral colonies (‘Colony’) of *Pocillopora verrucosa-Cladocopium pacificum* collected from control and heat aquariums (‘Treatment’) at different timepoints (‘Time (h)’). The total number of viral reads in each metatranscriptome is listed under ‘Viral reads’. Relative abundances of hits to each reference sequence are given as percentages with the total number of reads in parentheses. ‘Alignment metrics’ (mean e-values, bitscores and ID (%) ± standard deviation) were averaged across all reads that aligned to the given reference sequence within the five metatranscriptomes. ‘Mean bitscore’ and ‘Mean e-value’ indicate mean bitscores and e-values of the alignments, respectively. ‘Mean ID (%)’ indicates the percentage of characters that match exactly between the two sequences. SD: standard deviation.

| **A** | *Metatranscriptome* | | | *Reference sequence accession number* | | |
| --- | --- | --- | --- | --- | --- | --- |
|  |  |  |  | *RdRp* | *mcp* | |
| *Colony* | *Treatment* | *Time (h)* | *Viral reads* | *AOG17585.1* | *AOG17586.1* | *AOS87317.1* |
| 1 | Control | 0 | 78307 | 0.14 % (108) | 0.01 % (6) | 0.00 % (2) |
| 4 | Heat | 24 | 69405 | 0.54 % (369) | 0.04 % (24) | 0.04 % (22) |
| 5 | Control | 24 | 109612 | 3.15 % (3447) | 0.14 % (157) | 0.06 % (70) |
| 5 | Control | 108 | 46499 | 1.13 % (522) | 0.04 % (19) | 0.05 % (21) |
| 5 | Heat | 108 | 132709 | 0.94 % (1196) | 0.04 % (72) | 0.02 % (41) |
| **B** | *Alignment metrics* | | | *Reference sequence accession number* | | |
|  |  |  |  | *RdRp* | *mcp* | |
|  |  |  |  | *AOG17585.1* | *AOG17586.1* | *AOS87317.1* |
| Mean e-value ± SD | | | | 2.3·10^-5^ ± 9.9·10^-5^ | 5.5·10^-5^ ± 1.7·10^-4^ | 2.5·10^-5^ ± 1.2·10^-4^ |
| Mean bitscore ± SD | | | | 73.6 *±* 19.0 | 63.7 *±* 11.1 | 66.9 ± 12.8 |
| Mean ID (%) ± SD | | | | 60.1 *±* 7.3 | 58.4 *±* 9.3 | 53.3 ± 9.1 |

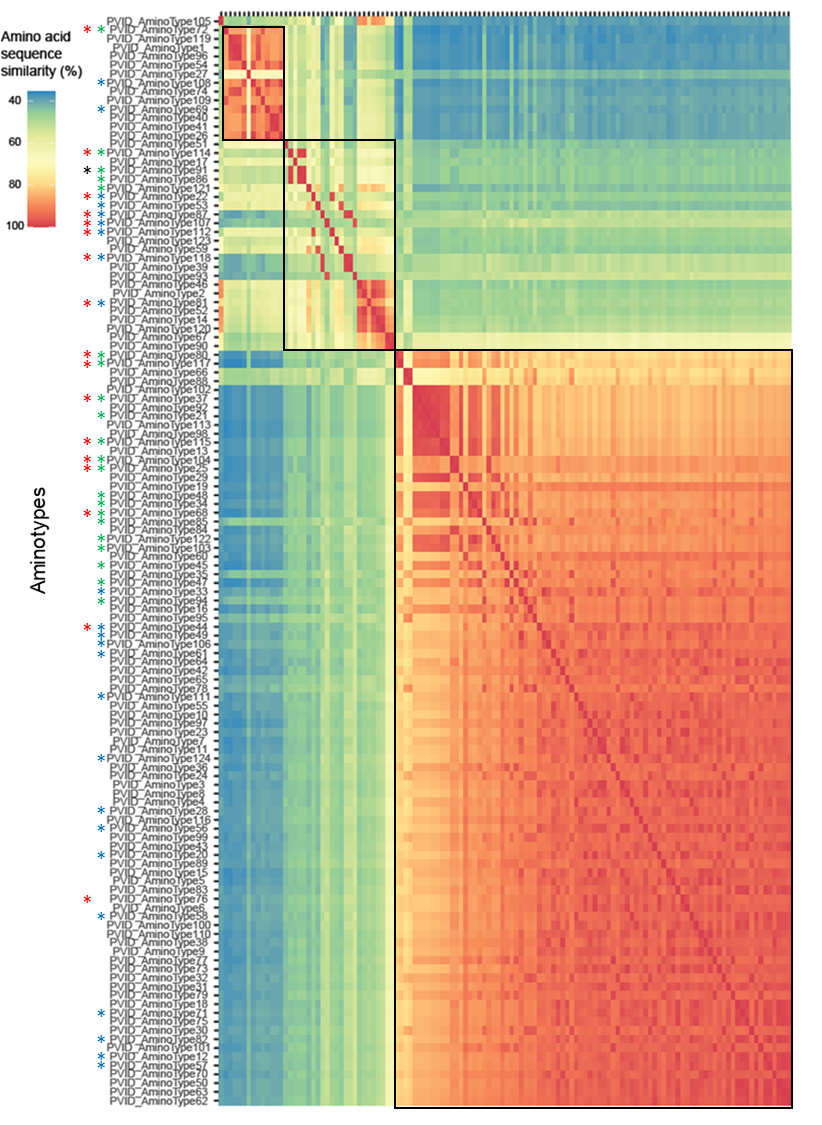

**Figure S2**	Similarity matrix (%) of all dinoRNAV major capsid protein (*mcp*) aminotypes (unique amino acid sequences) recovered from gene amplicon sequencing in this study. The three clusters of similar dinoRNAV aminotypes may represent individual RNA virus quasispecies. Blue asterisks indicate aminotypes unique to *Pocillopora verrucosa-Cladocopium pacificum*; green asterisks indicate aminotypes unique to *Pocillopora ligulata-Cladocopium latusorum.* Red asterisks indicate aminotypes unique to heat-treated fragments; black asterisks indicate aminotypes unique to control fragments.

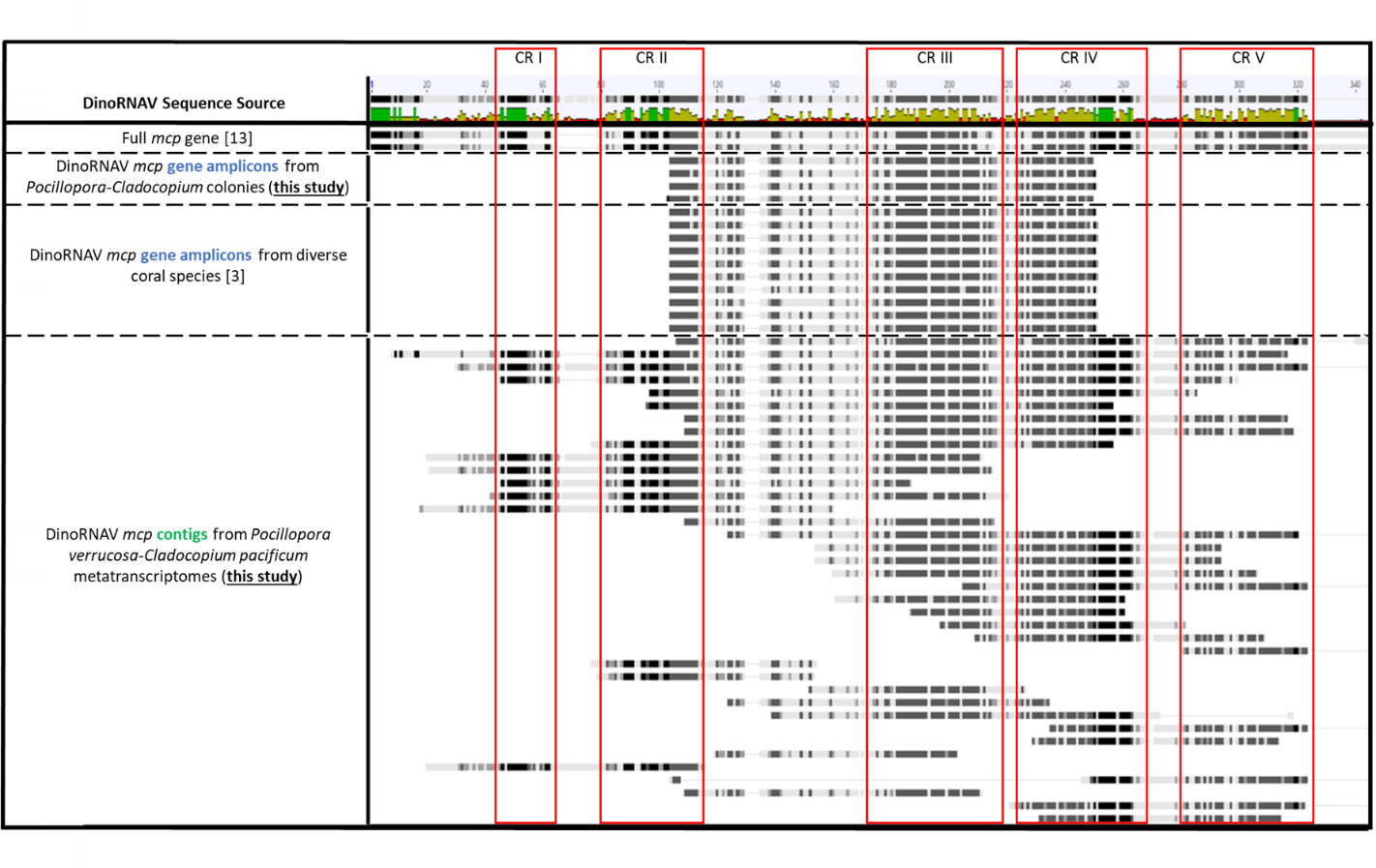

**Figure S3**	Mapping of dinoRNAV major capsid protein (*mcp*) amino acid sequences (this study) to the full *mcp* gene assembly from Levin et al. [13] (‘Full *mcp* gene’) reveals that the *mcp* gene sequences reported here are diverse but contain up to 5 putative conserved regions (CR I-V, red rectangles). There are also no visible large insertions or deletions in the sequences compared to the full genome. Together, these results indicate that the *mcp* gene amplicons and contigs reported here are from virions or viral infections, and are not endogenized viral elements. Sequences compared to the Levin et al. [13] full *mcp* gene assembly are from *Pocillopora-Cladocopium* colonies (*mcp* gene amplicon sequencing and contigs from metatranscriptomes; this study), as well as from a previous study analyzing dinoRNAV *mcp* gene amplicons from six coral species dominated by different Symbiodiniaceae [3].
